# Parametric Engineering of Atrioventricular Living Valve Transplants

**DOI:** 10.64898/2026.06.16.732756

**Authors:** Paul R. Kaneelil, Katrina J. Hon, Dominic P. Recco, Nikhil Thatte, Gianna Dafflisio, Peter E. Hammer, L. Mahadevan, Sitaram Emani

## Abstract

Diseases of the (mitral and tricuspid) atrioventricular valves (AVV), which regulate inflow from the atria to the ventricles, can result in severe obstruction to inflow (stenosis) or valvular leakage (regurgitation), requiring surgical intervention. In patients with small annulus diameters (**< 19** mm), valve replacement is a clinical challenge limited by prosthesis size constraints, lack of growth potential, suboptimal durability, and elevated thrombosis and bleeding risk. While living valve transplantation (LVT) has re-opened the possibility of using allogeneic valve tissue capable of growth and remodeling, translating this to the AVV has been challenging given the anatomical complexity of the sub-valvular apparatus. Here, we propose a strategy using a replacement bi-leaflet cylindrical valve fabricated from donor AVV tissue and artificial chordae, with a geometry designed to mimic the native AVV and engineered to satisfy predefined clinical targets. Pulse duplicator experiments allowed characterization of valve dynamics in terms of clinically important attributes framed as dimensionless parameters. A multi-objective optimization allowed us to identify an optimal design which we implemented in porcine AVV replacements (n=6). Our results demonstrated favorable hemodynamics with minimal regurgitation and stenosis, suggesting a promising method for patient-optimized valve replacements.

---

The atrioventricular valves (AVV), including mitral and tricuspid valves, regulate inflow from the atria into the ventricles. Disease of the AVV can result in obstruction to inflow (stenosis) or valvular leakage (regurgitation) [1]. Although AVV repair is preferred when feasible, replacement remains necessary in cases of irreparable valve pathology, with over 19,000 atrioventricular valve replacements (AVVR) performed annually in the U.S.[1–3]. Outcomes of AVVR in pediatric patients are compromised by the need for reoperation, with more than half of the children requiring repeat AVVR within 10 years, most commonly due to a patient outgrowing the valve or progressive deterioration of the valve materials [4, 5]. These challenges are particularly pronounced in patients with small annulus diameters (< 19 mm), and no valve prostheses are available for diameter less than 15 mm [4, 6–8]. Among neonates undergoing AVVR, reoperation rates approach 50% within the first year [9–11].

Current valve replacement options include bioprosthetic valves composed of bovine or porcine tissue, or mechanical valves constructed from pyrolytic carbon [12]. Both approaches have important limitations. Bioprosthetic valves consist of nonliving tissue susceptible to microthrombosis and scarring, resulting in early structural degeneration with approximately 80% demonstrating dysfunction or requiring reoperation by 10 years [13]. The rigid scaffolding structure that houses the valve additionally reduces the effective valve orifice area, predisposing to stenosis. Mechanical valves require lifelong anticoagulation and are associated with approximately 20% higher risk of bleeding and thrombosis compared to bioprosthetic valves [13–15]. Critically, neither prosthesis type possesses growth potential, resulting in the need for repeated interventions as pediatric patients outgrow the valve [2, 16, 17]. Consequently, patients with small annuli experience substantial morbidity and mortality related to residual valve disease and multiple reinterventions [18, 19].

Living valve transplant (LVT) is an alternative strategy that provides potential for growth, long term durability, and resistance to thrombosis. [20, 21]. This approach involves the transplantation of only the valvular component of the freshly procured donor heart and provides allogeneic valve tissue with viable cellular components, capable of growth and self-repair. Early experience with aortic and pulmonary LVT demonstrates favorable hemodynamic performance and normal appearance of valves by echocardiography. [22, 23]. However, extending LVT to the AVV position remains challenging given the anatomical complexity of the sub-valvular apparatus. The AVV possesses a geometrically complex apparatus consisting of a multi-leaflet conical structure [24] supported by a fibrous annulus and a network of chordae, as shown in Fig. 1(a.i)-(b.i). We hypothesize that replicating key features of the native AVV geometry in a replacement valve will lead to favorable hemodynamic performance. In this work, we introduce the Live-AVV, a bi-leaflet biosynthetic AVV that approximates native AVV geometry, constructed from living donor valve tissue and expanded-polytetrafluoroethylene (ePTFE) chordae, as schematically represented in Fig. 1(b.ii), The design is optimized to meet predefined clinical targets and is scalable to fit individual patient anatomy. This approach enables a standardized patient-specific LVT strategy that avoids donor size constraints and complications associated with transplantation of donor subvalvular structures, including technical complexity, ischemia, and dehiscence.

**Fig. 1.**
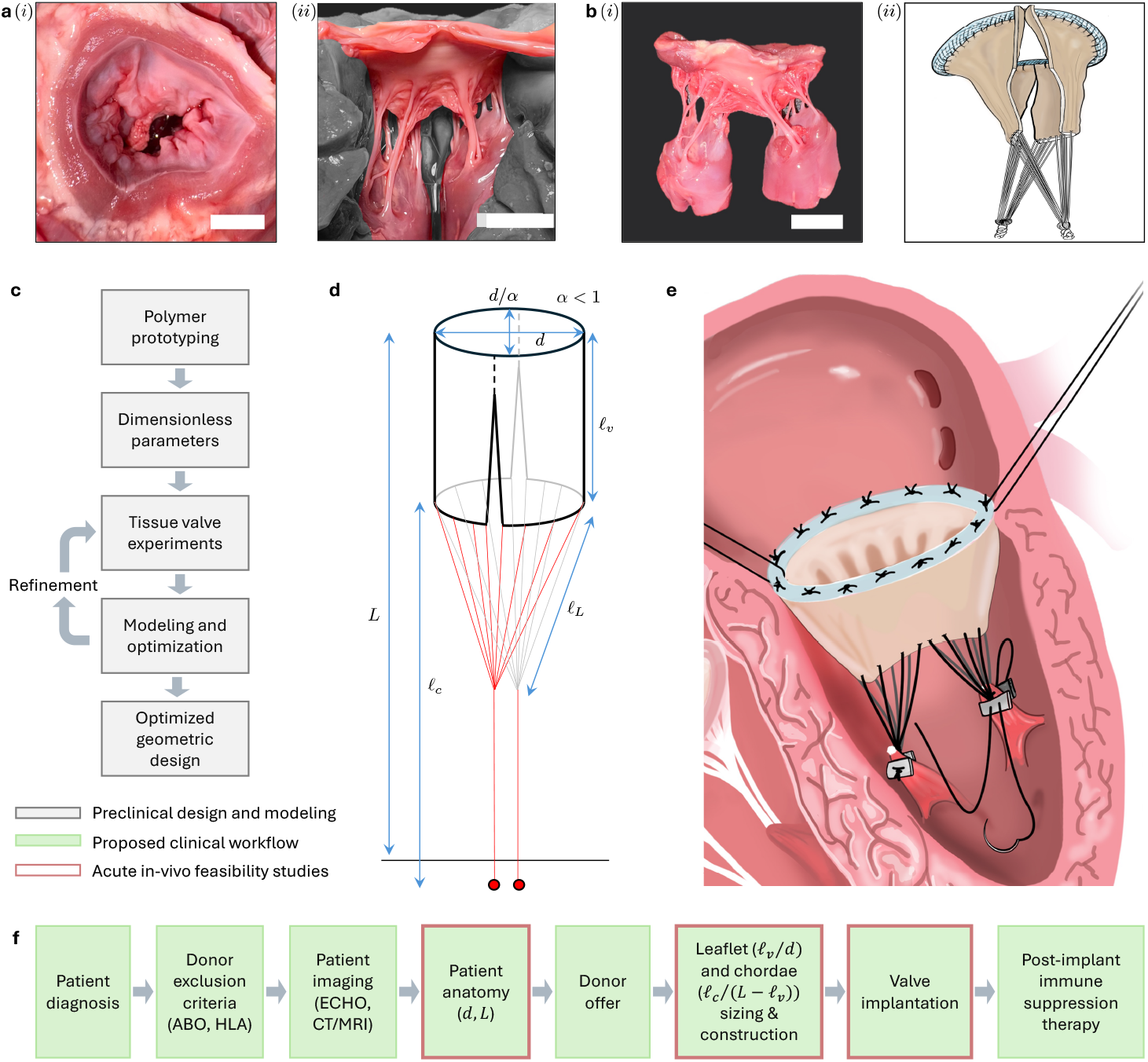
Atrioventricular valve (AVV) geometry and transplant strategy. (a.i) Porcine mitral valve (MV) in an excised heart from (i) Surgeon’s view and (ii) side view showing the complex chordae tendineae and papillary muscle structure. (b) (i) An excised MV apparatus showing the leaflets, chordae, and papillary muscles. All scale bars represent 10 mm. (ii) Schematic of an engineered AVV. (c) Experimental workflow for in-vitro tissue valve experiments and modeling used to optimize Live-AVV geometry. (d) Schematic of the geometry of the bi-leaflet cylindrical Live-AVV with an elliptic annulus, illustrating key geometric parameters derived from polymer prototype studies. (e) Schematic of the Live-AVV constructed from donor valve tissue leaflets and ePTFE chordae and implanted into a recipient heart. (f) Envisioned surgical workflow for clinical translation, including key surgical steps included for acute in-vivo feasibility studies (blue outline).

To achieve this, we identified the key geometric and structural features required to engineer a functional Live-AVV. Figure 1(c),(f) illustrates the workflow, including the preclinical design optimization stage, in-vivo feasibility studies, and an anticipated clinical translation workflow. A crucial step was determining the size of the leaflets and length of the chordae such that the constructed Live-AVV met the functional standards of a normal AVV. Building on our previous work characterizing the closing dynamics of a conical valve using a polymeric prototype [25], we incorporated chordae and commissural slits to create a bi-leaflet valve and identified the minimal set of governing dimensionless parameters (See the Supplementary Information). These parameters were then systematically varied and evaluated in-vitro under physiological conditions in a pulse duplicator, enabling identification of geometries that satisfied clinically relevant hemodynamic criteria. The optimized design was subsequently applied to Live-AVV (n=6) implantation into tricuspid valve position in an acute large animal model, demonstrating favorable hemodynamic performance. Collectively, these studies established a standardized framework for constructing a geometrically optimized Live-AVV engineered to reproduce the functional performance of a native AVV.

## 1 Results and Discussion

### 1.1 Live-AVV geometry

For the design of the Live-AVV, we identified the minimal form that was required to maintain the function of the AVV. A native AVV, as shown in Fig. 1(a)-(b), consists of two main leaflets attached at the valve annulus, with numerous chordae tendineae connecting the free edge and the ventricular surface of the leaflets to the papillary muscles of the ventricle. [24, 26]. This complex geometry can be characterized by several parameters - annulus diameter, the annulus eccentricity, leaflet length and width, leaflet shape, chordae length, number of chordae, chordae attachment sites and angles, and slit length, defined as the length of commissural opening between the two leaflets.

We performed experiments with a polymeric conical valve, as explained in detail in the Supplementary Information, to determine the important geometric parameters. In diastole when the valve is open, the AVV resembles the shape of a funnel[26] or a cone [24], due in part to the angle of the supporting chordae. Experiments with polymeric conical valve showed that the length of the valve *ℓ*_*v*_ and the effective chordae length *ℓ*_*c*_ were important parameters that impacted valve leakage during systole with larger *ℓ*_*v*_ and intermediate *ℓ*_*c*_ associated with minimal leakage. In contrast, the slit length and the opening diameter impacted valve performance during diastole (stenosis) with larger values of both associated with minimal stenosis. For the Live-AVV, slit length was maximized to minimize stenosis as much as possible. In contrast to most commercially available prosthetic valves which exhibit transvalvular gradients of 5-6 mmHg, we anticipated that this design, with maximal slit length and without the geometric constraints of a tubular structure, would produce lower transvalvular gradients [27–29].

Based on these results from the polymeric valve experiments, we found that the relevant variables can be organized into three dimensionless geometric parameters that define the geometry and control the function of the valve: a rescaled valve length *ℓ*_*v*_*/d*, a rescaled chordae length *ℓ*_*c*_*/*(*L* − *ℓ*_*v*_), and an anatomic aspect ratio *L/d*. A schematic of the geometry of the Live-AVV, a bi-leaflet elliptical cylinder, is shown in Fig. 1(d). Here, the smaller diameter of the ellipse is *d*, the ratio of minor-to-major diameter is *α*, and the length from the annulus to the papillary attachment location is *L*. The goal of the in-vitro experiments was to determine the optimal values of *ℓ*_*v*_*/d* and *ℓ*_*c*_*/*(*L - ℓ*_*v*_), given *L/d*. We did not expect the results to be a strong function of *L/d* as long as *L/d* ≫ *ℓ*_*v*_*/d*; results from experiments varying *L/d* are summarized in the Supplementary Information.

### 1.2 In-vitro experiments

We constructed Live-AVVs utilizing porcine tricuspid valve tissue sutured onto a polymer annulus (Vinylpolysiloxane; Zhermack 32 shore A) to perform in-vitro experiments in a pulse duplicator (ViVitro Labs; Victoria, BC, Canada). Five different geometries were tested (n=3 each): *ℓ*_*v*_*/d* = 0.625, 0.75, 0.875, 1.0, 1.25. Figure 2(a) shows an image of the constructed Live-AVV. For each geometry, we varied the dimensionless chordae length, or *ℓ*_*c*_*/*(*L−ℓ*_*v*_), in the pulse duplicator. The following parameters were fixed for all geometries: *α* = 0.9, *ℓ*_*L*_ = 20 mm, d = 14.4 mm, and L = 47 mm (*L/d* = 3.26). (See the Methods for more details). Note that for ease of in-vitro testing, *ℓ*_*L*_ was fixed while changing *ℓ*_*c*_. The effect of varying *ℓ*_*L*_ is discussed in the Supplementary Information. In addition, the number of chordae (8 per leaflet) was kept contant for all geometries. Considering a peak systolic pressure of 120 mmHg, 8 chordae per leaflet results in approximately 0.18 N per chordal loop which is more than an order of magnitude less than the average knot pull tensile strength of 1.35 N of a CV-7 ePTFE suture [30].

**Fig. 2.**
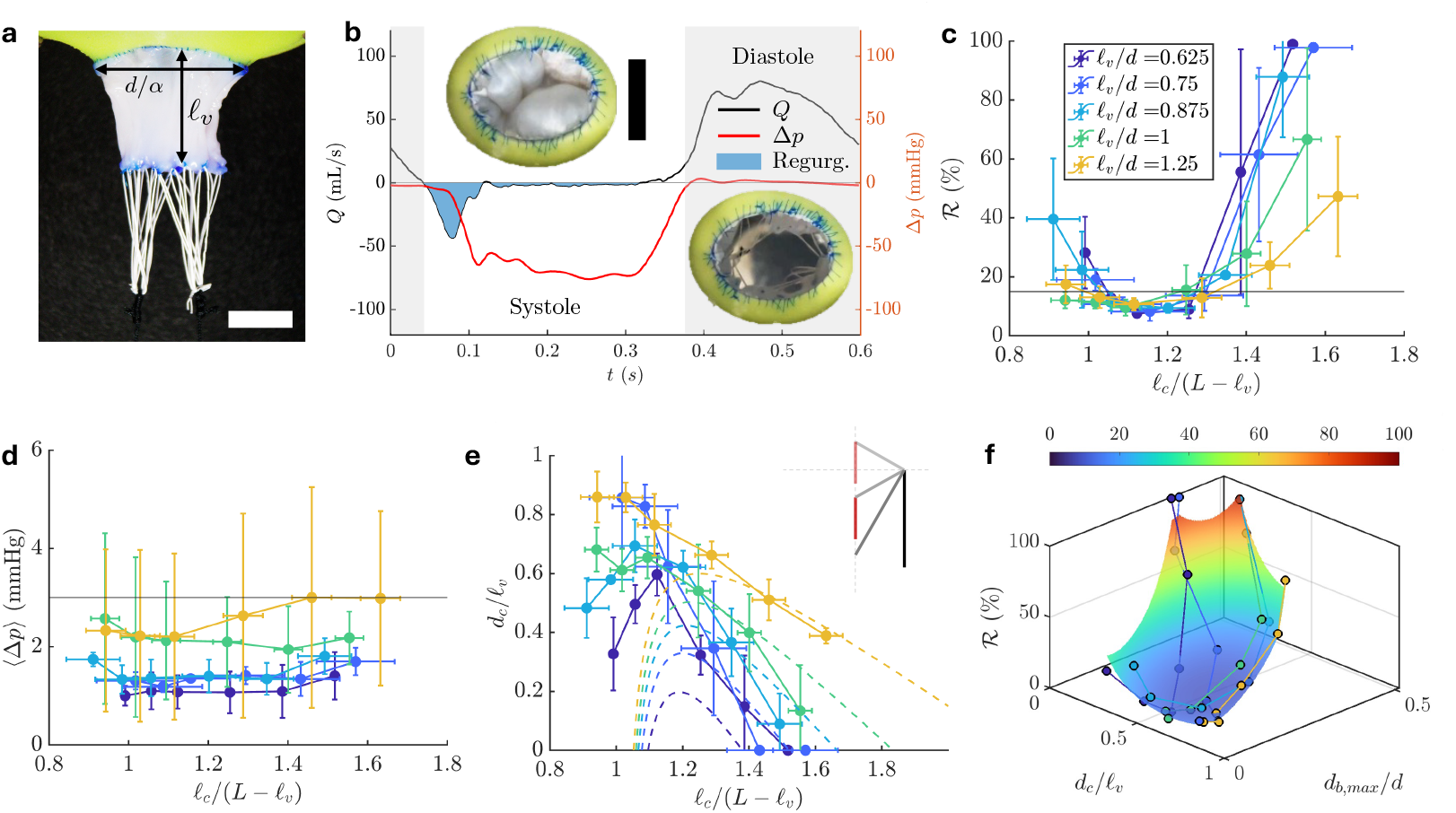
In-vitro testing of Live-AVVs. (a) Image of a Live-AVV constructed from porcine tricuspid tissue supported by ePTFE chordae. The Live-AVV are sutured on to a polymer annulus of pre-determined size (green). (b) Pressure data from the pulse duplicator showing the transvalvular pressure (Δ*p*; solid red line) averaged over ten cardiac cycles. The flow rate *Q* across the valve (black line) shows the total regurgitation volume shaded blue. Note that the positive flow correspond to diastole (gray region) and negative flow correspond to systole (white region). The insets show representative surgeon’s view images during systole and diastole. (c) The total regurgitant fraction as a function of the dimensionless chordae length shows a range where ℛ falls below the clinical target of 15% (black horizontal line). Note that the lines connecting the data points are a visual guide. The legend indicates the different colors corresponding to the *ℓ*_*v*_ */d* values; this same legend applies wherever these colors appear throughout the figure. (d) The mean transvalvular gradient as a function of the dimensionless chordae length, where all the mean data is below the clinical target of 3 mmHg. (e) The rescaled coaptation height *d*_*c*_*/ℓ*_*v*_ as a function of dimensionless chordae length. The dashed lines are the predictions from the geometric model in Eq. (1). (f) The regurgitant fraction as a function of *d*_*c*_*/ℓ*_*v*_ and *d*_*b,max*_*/d* showing their complex non-linear relationship. The surface is from an exponential regression. All scale bars represent 10 mm.

The in-vitro experiments in the pulse duplicator were performed under physiological conditions corresponding to the annulus size of the Live-AVV. See the Methods section for specific flow conditions. A typical transvalvular pressure (Δ*p*; red line) and flow rate (*Q*; black line) profile during one cardiac cycle is shown in Fig. 2(b). Positive flow rate corresponds to diastole which is shaded gray and negative flow rate corresponds to systole. The insets show surgeon’s view images of the valve in both diastole and systole, where it is seen to closely resemble the morphology of a native valve. The area corresponding to the total regurgitation volume is shaded in light blue in Fig. 2(b).

The total regurgitant fraction *R* was defined as the ratio of regurgitation volume to stroke volume. Figure 2(c) shows the regurgitant fraction as a function of *ℓ*_*c*_*/*(*L* − *ℓ*_*v*_) for various leaflet geometries *ℓ*_*v*_*/d*. The black line denotes the clinical target of *R* < 15% [31], consistent with clinical definitions of normal to mild regurgitation and established International Organization for Standardization (ISO) device performance standards [32–34]. Note that three replicates were made for each Live-AVV geometry and the error bars represent the standard deviation from the three tests. The regurgitant fraction increases when chordae length is greater than about *ℓ*_*c*_*/*(*L* − *ℓ*_*v*_) ‘1.3, due to inadequate leaflet coaptation and incomplete valve closure of the valve leaflets. As chordae length decreases, we observe a range of *ℓ*_*c*_*/*(*L*−*ℓ*_*v*_) within which the *R* is below the clinical target. Further decrease in chordae length *ℓ*_*c*_*/*(*L* − *ℓ*_*v*_) ⪅ 1 is associated with increase in *R* due to leaflet tethering. Notice that *ℓ*_*c*_*/*(*L* − *ℓ*_*v*_) = 1 represents the case in which the effective chordae length is equal to the distance between the attachment point and the edge of the free standing Live-AVV leaflets, corresponding to a taut chordae during diastole. Thus, values of *ℓ*_*c*_*/*(*L* − *ℓ*_*v*_) < 1 represent the case in which the leaflets are stretched beyond resting dimensions and *ℓ*_*c*_*/*(*L* − *ℓ*_*v*_) > 1 represent the case in which there is redundant chordae length during diastole. For larger *ℓ*_*v*_*/d*, the curvature of the regurgitation data is reduced, leading to a smaller deviation from the clinical target across *ℓ*_*c*_*/*(*L* − *ℓ*_*v*_). In comparison, smaller *ℓ*_*v*_*/d* exhibit pronounced sensitivity, with *R* increasing by nearly six-fold with modest changes in *ℓ*_*c*_*/*(*L* − *ℓ*_*v*_). These findings suggest that larger leaflet geometries provide greater tolerance to variations in chordae length, enhancing design robustness and maintaining near-optimal performance across a broader parameter range. At high *R*, small changes in *ℓ*_*c*_*/*(*L* − *ℓ*_*v*_) produce larger, more variable reductions in regurgitation, reflected by both larger error bars and greater stepwise changes between adjacent conditions, whereas near the clinical threshold, similar adjustments yield only incremental gains. The mean transvalvular pressure ⟨Δ*p*⟩ is defined as the mean gradient during diastole, Δ*p*> 0, and is a parameter that quantifies stenosis. Figure 2(d) shows 6.*p* from our in-vitro experiments, with the black line representing the a maximum acceptable pressure gradient of 3 mmHg, selected as a conservative benchmark that sits just above normal AVV hemodynamics (≤ 2 mmHg) and remains within the threshold for mild stenosis [31, 35]. All of the mean data is below this clinical target. We did not observe a strong trend in ⟨Δ*p*⟩ as a function of *ℓ*_*c*_*/*(*L* − *ℓ*_*v*_), but it increases non-linearly with increasing *ℓ*_*v*_*/d* as expected considering an approximate Poiseuille type flow, where Δ*p* ~ *ℓ*_*v*_*/d*^4^. Thus, the benefit of greater tolerance in ℝ observed with longer leaflet lengths is counteracted by an increase in ⟨Δ*p*⟩.

In addition to the hydrodynamic parameters, we also quantified, via echocardiography, the coaptation height *d*_*c*_, which is the length of leaflet contact or coaptation in systole, and the maximum billow height *d*_*b,max*_, which is the maximum displacement of the leaflet into the atrium in systole, referenced from the annulus plane. Both of these parameters are of clinical significance as they have been associated with long term durability of the valve apparatus [36–42]. See the Methods section for more details regarding imaging and measurement and see the Supplementary Information for the raw data for *d*_*c*_ and *d*_*b,max*_. The rescaled coaptation height *d*_*c*_*/ℓ*_*v*_ for all the experiments are plotted in Fig. 2(e) and has an inverted U-shaped dependence on *ℓ*_*c*_/(*L − ℓ*_*v*_). As *ℓ*_*c*_*/*(*L − ℓ*_*v*_) decreases from the upper extreme when the Live-AVV is everted without any coaptation, *d*_*c*_/*ℓ*_*v*_ increases to reach a maximum in most cases before decreasing again. Intuitively, this decrease corresponds to the cases where the chordae are too short and cause lateral tethering of the leaflets, restricting them from coapting centrally. The Live-AVVs with larger *ℓ*_*v*_*/d* do not show a significant decrease in *d*_*c*_/*ℓ*_*v*_, presumably due to the abundance of leaflet material that allows for stretching. Leaflet stretching plays a significant role in providing ample coaptation. In Fig. 2(e), the dashed lines correspond to a geometric model of coaptation given by the dimensionless equation,

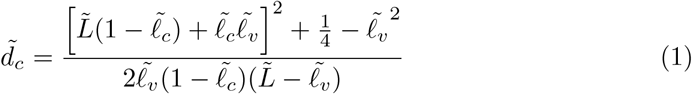

Where 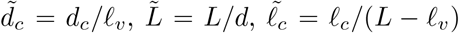, and 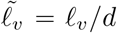. Because the geometric model neglects stresses arising from elasticity and fluid pressure, we expect it to underestimate coaptation height for short chordae, where stretching dominates, and overestimate it for long chordae, where bending dominates. In Fig. 2(e), we see that the experimental measurement of *d*_*c*_/*ℓ*_*v*_ is always larger than the geometric prediction for small *ℓ*_*c*_/(*L* − *ℓ*_*v*_) and that it is smaller than the geometric prediction for large *ℓ*_*c*_/(*L* − *ℓ*_*v*_).

We expect the regurgitant fraction of the AVV to be strongly coupled to the coaptation and billow heights [36–42]. We therefore examined the relationship between *R*, coaptation and billowing height in Live AVV. Figure 2(f) shows *R* as a function of *d*_*c*_/*ℓ*_*v*_ and *d*_*b,max*_*/d*. Note that the relationship between these parameters is nonlinear. To quantify the relationship, we performed an exponential regression (*R*^2^ = 0.88) of the form log(*R*) *~ β*_1_(*d*_*c*_/*ℓ*_*v*_)+ *β*_2_(*d*_*b*_*/d*)+ *β*_3_(*d*_*c*_/*ℓ*_*v*_)^2^ + *β*_4_(*d*_*b*_*/d*)^2^ + *β*_5_(*d*_*c*_/*ℓ*_*v*_)(*d*_*b*_*/d*). The minimum of this surface occurs at *d*_*c*_*/ℓ*_*v*_ = 0.59 and *d*_*b,max*_*/d* = 0.17. Unlike a native valve in which billowing is often associated with reduced coaptation height and therefore higher regurgitation, this result suggests that the Live-AVV reaches maximum coaptation and minimum regurgitation when there is a small amount of billow. This is presumably a consequence of the lack of secondary chordae in our design.

Additional experiments on Live-AVV were performed to evaluate the effects of varying some of the parameters that were initially held constant for the in-vitro experiments. We found that loop lengths *ℓ*_*L*_ ≥ *d/*2 demonstrated comparable regurgitant fractions and transvalvular pressure gradients to one another, while reducing slit length by half reduced regurgitant fraction at extreme chordae lengths, though increased the risk for stenosis with smaller valve lengths *ℓ*_*v*_*/d*. Full methodological details and raw data are provided in the Supplementary Information.

## 1.3 Design optimization

We used the results from the in-vitro experiments to determine the clinically feasible parameter region in our design space and to further determine the optimal values of *ℓ*_*v*_*/d* and *ℓ*_*c*_*/*(*L − ℓ*_*v*_) for our given *L/d*. Figure 3(a-d) shows the experimental data (black dots) for the four clinical measures fitted to a surface (colormap) using a Gaussian Process Regression (GPR) as a function of *ℓ*_*v*_*/d* and *ℓ*_*c*_/(*L − ℓ*_*v*_). This fit (*R*^2^ > 0.95) captures the smooth nonlinear dependencies without imposing a specific parametric form on the complex form-function relationship of the Live-AVV. The clinical targets for these measures are indicated with the black contour lines. The clinical target for *d*_*c*_ is 5 mm based on normal AVV coaptation height [37] and the clinical target for *d*_*b,max*_ is 3 mm. Although physiological leaflet excursion is often defined as ≤ 2 mm, a slightly higher threshold was adopted to account for the absence of secondary chordae in this model [43].

**Fig. 3.**
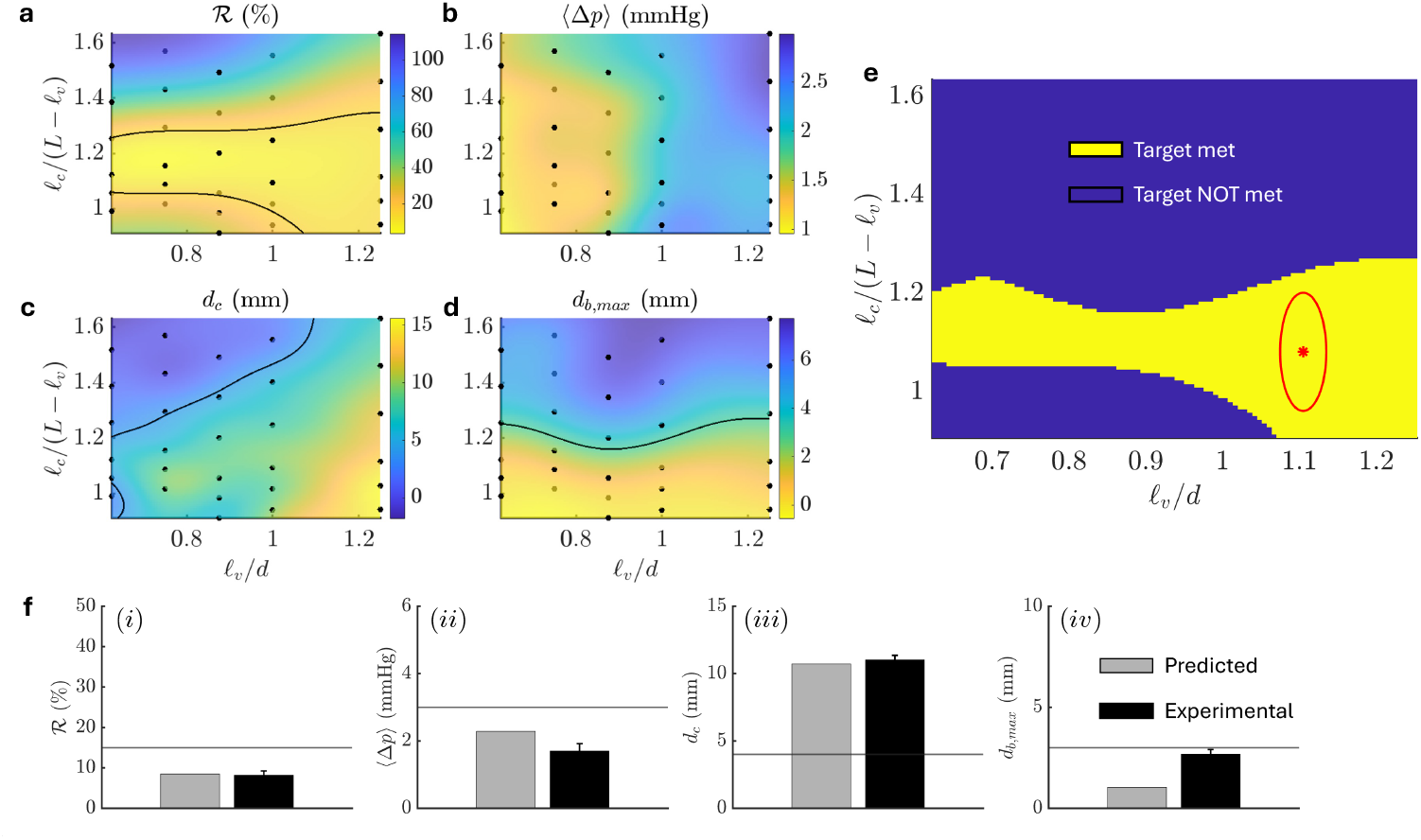
Geometrical design space and optimization. Data (black dots) for the four clinical measures,, *R* (b) ⟨Δ*p*⟩, (c) *d*_*c*_, and (d) *d*_*b,max*_, are shown as a function of *ℓ*_*v*_*/d* and *ℓ*_*c*_*/*(*L* − *ℓ*_*v*_). The colormap represents a surface fit using Gaussian Process Regression (GPR) and the black lines are contours along the corresponding clinical targets. (e) Combined design space of the Live-AVV for *L/d* = 3.26, based on fixed Live-AVV parameters. Here, the yellow represents the region where all clinical targets are met. The red star marks the optimal design according to our optimization framework, while the red ellipse denotes the considered potential variability in heart geometry over a cardiac cycle (including a safety factor), which still lies entirely within the target-satisfying region. (f) Results of the optimally designed tissue Live-AVVs from in-vitro experiments. The solid black bars correspond to the experiment and the gray bars correspond to the predicted values from the GPR models.

In Fig. 3(e), we show the combined design space in binary form with blue representing the region where the clinical target is not met and yellow representing the region where all the targets are met. While any combination of *ℓ*_*v*_*/d* and *ℓ*_*c*_/(*L − ℓ*_*v*_) in this region should produce a clinically acceptable Live-AVV, we performed a multi-objective optimization on the data to derive a more *optimal* design criterion.

The first trivial step in the optimization essentially determines the region where all clinical targets are met – i.e. the yellow region in Fig. 3(e) – and thereby addresses dynamic valve function. In the second step, we impose three additional constraints. First, the optimal criterion must account for dynamic heart function, i.e., physiological variation in ventricular geometry of approximately *±*4% *ℓ*_*v*_/*d* [44] and *±*7.4% *ℓ*_*c*_/(*L − ℓ*_*v*_) [45] throughout the cardiac cycle, such that it is surrounded by an elliptical neighborhood of the corresponding size where all targets remain satisfied. We add a safety factor of 1.5 to these values to be more conservative, and to also account for variability in mechanical properties of the valve (See Supplementary Information). Second, we account for tissue availability by minimizing *ℓ*_*v*_/*d* and therefore favor designs that require less donor tissue. In practice, donor valve size represents a significant constrant and is partly driven by our rectangular leaflet design, which demands more tissue than the native AVV’s lobed shape, characterized by minimal commissural and maximal central leaflet length. Future work may focus on developing a more tissue-conservative design in addition to evaluating performance of composite versus single-piece leaflets. Third, we penalize sensitivity to regurgitation by minimizing the curvature of *R* vs. *ℓ*_*c*_/(*L* − *ℓ*_*v*_) such that there is a broad basin of acceptable performance. These latter two constraints are both weighed equally at 0.75. Under this framework, the optimal design is *ℓ*_*v*_/*d* = *ℓ*_*c*_/(*L* − *ℓ*_*v*_) = 1.1 (red star in Fig. 3(e)), with the ellipse indicating the accounted variation in ventricular geometry.

To verify the reliability of this optimality result, we constructed three Live-AVVs of *ℓ*_*v*_/*d* = *ℓ*_*c*_/(*L* − *ℓ*_*v*_) = 1.1 and tested them in-vitro. The clinical measures of these optimized Live-AVVs are summarized in Fig. 3(f). We note that clinical thresholds for all four clinical targets were met, with *R*, ⟨Δ*p*⟩, and *d*_*c*_ performing better than predicted. Experimental *d*_*b,max*_ (2.7 mm) was slightly higher than predicted (1.0 mm), although it remained within clinical targets (see Fig. 3(f)). This difference may reflect variability in tissue mechanical properties.

### 1.4 In-vivo experiments

We performed six prototype Live-AVV implants in a porcine model of AVV replacement (n=3 Yorkshire pigs, *L/d* ∈ [1.54, 2.46]), as described in detail in the Methods section. Live-AVVs were constructed from donor human tricuspid valve tissue and implanted in the tricuspid position of 6-8 kg female Yorkshire pigs. The tricuspid position was selected for initial studies to enable evaluation in the beating heart without the need for cardiac arrest and lower-pressure afterload conditions. Two implantation strategies were evaluated. In the ringed configuration Live-AVV was constructed within a polymer ring and subsequently the ring was sutured into the native tricuspid annulus after native leaflet resection as shown in Fig. 4(a),. This configuration replicated the in-vitro configuration and enabled preoperative assembly. In the non-ringed configuration, leaflets were sutured directly to the native annulus, as shown in Fig. 4(b). Full operative details and implantation sequence are provided in the Methods section. Notably, a larger annulus size in the third animal necessitated composite leaflet construction. Figure 4(c-d) shows representative epicardial echocardiogram images of the implanted Live-AVV in diastole and systole from a parasternal long-axis view. Echocardiographic assessment was performed following complete weaning from cardiopulmonary bypass (CPB), when feasible. In 1/6 Live-AVV, separation from CPB was not feasible due to poor cardiac contractility. Valvular assessment for this configuration was therefore obtained on partial bypass support at a flow of 1 L/(min · m^2^). A total of *n* = 3 ringed and *n* = 3 non-ringed Live-AVV were completed in this study.

**Fig. 4.**
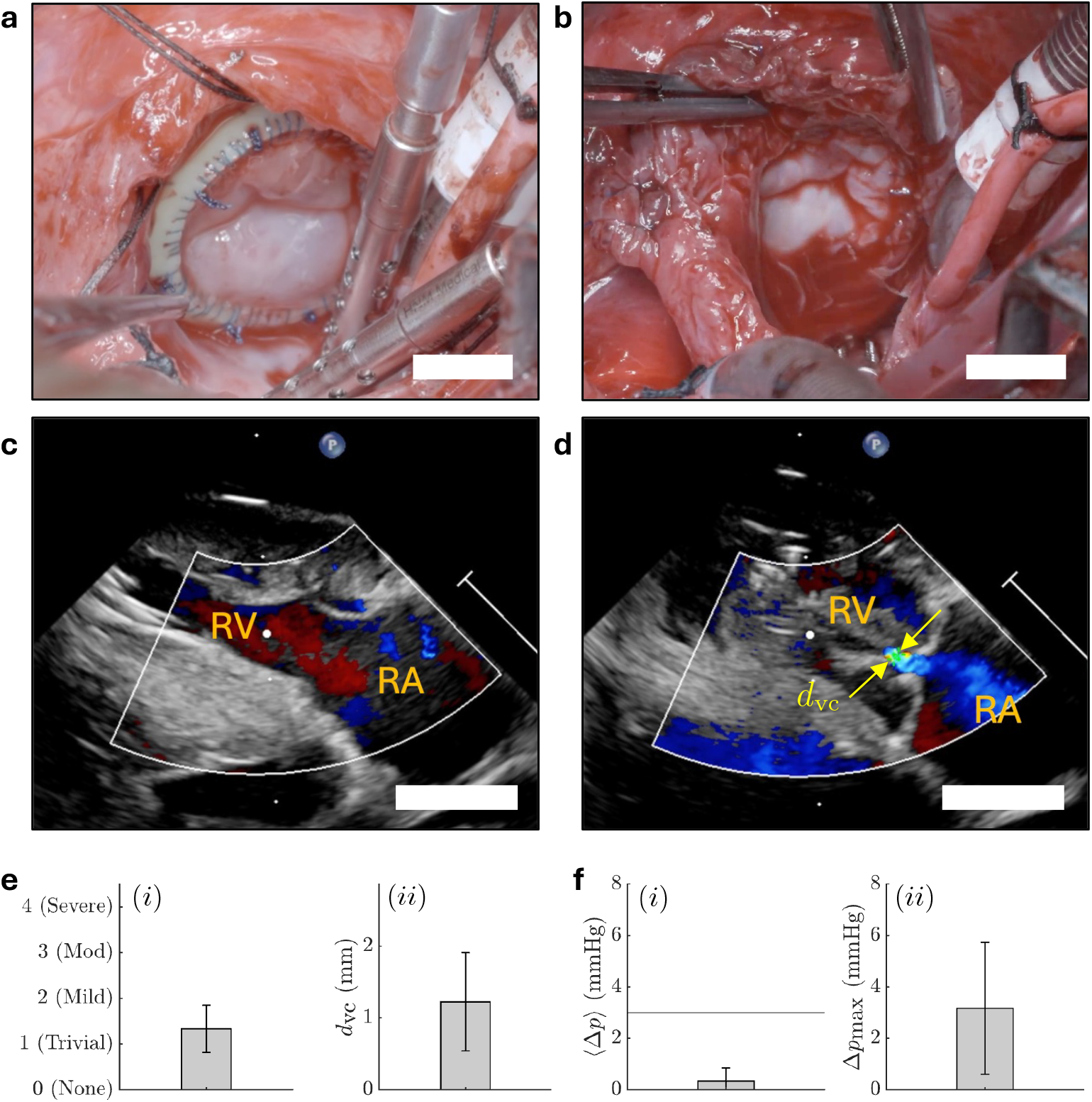
In-vivo testing of Live-AVVs. (a) Implanted Live-AVV sutured onto a polymeric annulus ring (green). (b) Implanted Live-AVV sutured directly onto the native annulus (no ring). (c-d) Parasternal long-axis echocardiographic views of an implanted Live-AVV in diastole and systole, respectively. Regurgitation was further quantified by measuring vena contracta diameter, *d*_vc_, for central jets (yellow arrows). All scale bars represent 10 mm. (e) Averaged regurgitation data from the in-vivo experiments showing (i) severity of regurgitation and (ii) *d*_vc_ across all in-vivo implants (*n* = 6). (f) Averaged pressure data showing (i) the mean transvalvular pressure ⟨Δ*p*⟩ well below the clinical target of 3 mmHg (black horizontal line) and (ii) the maximum transvalvular pressure Δ*p*_max_ from the in-vivo experiments.

Intraoperative epicardial echocardiography revealed trivial-to-mild regurgitation and no evidence of stenosis across all implanted Live-AVVs as shown in Fig. 4 (e)-(f). Regurgitation was further quantified by vena contracta diameter, *d*_vc_, a standard echocardiographic measure of regurgitant jet severity that reflects the size of the regurgitant jet, as shown in Fig. 4(d). Central jets ranged from non-measurable vena contracta with trivial flow to a maximum vena contracta diameter of 2 mm (mean 1.2 mm). In two implants, regurgitation was attributable to perivalvular leak, or leak at the annulus attachment point, suggesting a surgical implantation error which was immediately rectified with an additional reinforcing suture. Severe right ventricular dysfunction was present in one animal; despite this, valve function remained preserved, suggesting relative robustness of the Live-AVV geometry under impaired ventricular function.

Transvalvular gradients were measured by transthoracic echocardiography (TTE) Doppler images, with *mean* transvalvular gradients ranging from 0-1 mmHg (mean 0.3 mmHg). These findings were corroborated by invasive pressure measurements, which demonstrated *peak* transvalvular gradients ranging from 1-7 mmHg (mean 3.2 mmHg). Coaptation and billow height were quantified from imaging using the same methods as in-vitro experiments, with a mean coaptation height of 7.5 mm and a mean billow height of 0 mm across all implants. No qualitative differences in valve performance were observed between ring and no ring implantation strategies. Similarly, composite leaflet construction performed comparably to non-composite Live-AVVs.

Collectively, these data indicate that the optimized geometric configuration maintained Live-AVV competence with minimal transvalvular obstruction under physiological in-vivo loading conditions, meeting all clinical targets previously outlined. These findings demonstrate the feasibility of valve design that mimics native AVV geometry and provides a foundation for future evaluation in chronic studies. Several limitations should be acknowledged. The acute study design and narrow animal size range precluded assessment of long-term durability, tissue remodeling, growth, and scalability across the broad spectrum of patient anatomies encountered clinically. In-vivo evaluation was limited to the tricuspid position and did not assess valve performance under the higher-pressure loading conditions of the mitral position. Furthermore, the study design did not permit evaluation of host immune responses to allogeneic valve tissue, which will be important for defining rejection risk and immunosuppression requirements. Beyond AVV replacement, the geometric design principles described here may also inform leaflet augmentation or other reconstructive procedures for complex AVV disease. Future chronic studies in both AVV positions will be necessary to evaluate long-term valve function, host-graft interactions, growth potential, and scalability. Taken together, these findings establish a foundation for the development of living, geometrically optimized AVV replacement therapy for patients of all annulus sizes.

## 2 Conclusions

Our engineered Live-AVV extends the LVT paradigm from the semilunar valves to the AVV position. The design principles and methodology presented here provide a framework for constructing a custom AVV from living AVV tissue, with geometry tuned to meet clinically relevant targets. In-vitro experiments informed the optimal design of the valve, which was then successfully implemented in porcine AVV transplants. While our design demonstrated robust hemodynamic performance, this study was limited by its acute design and use of cryopreserved donor valve tissue, precluding assessment of growth, durability, and immune response. In addition, in vivo evaluation was restricted to the tricuspid position and therefore did not assess valve performance under the higher-pressure loading conditions of the mitral circulation. Next steps will incorporate living donor valve tissue with minimal ischemic time in chronic models and extend to the mitral position to evaluate leaflet adaptation over time, host-graft interactions, and the role of immunosuppressive therapy in supporting long-term valve function. Nevertheless, this work establishes a geometry-guided framework for engineering a Live-AVV and defines optimized design parameters as an initial step towards creating a living AVV that mimics native valve structure with potential for growth and durability. These findings provide a foundation for future chronic transplantation studies and, ultimately, clinical translation, offering a promising AVV replacement alternative for patients with limited treatment options.

## 3 Methods

### 3.1 In-vitro experiments: Tissue sourcing and preparation

To conduct primary in-vitro experiments, porcine tricuspid valves were harvested from hearts obtained from LAMPIRE Biological Laboratories, Inc. (Pipersville, PA, USA). Valve leaflets were dissected to remove excess ventricular and subvalvular attachments while preserving native leaflet architecture. Between dissection and Live-AVV construction, tissue was stored at −80°C to limit cellular degradation. During construction, leaflet tissue was maintained using hydrated physiological solution to prevent dehydration and preserve mechanical compliance.

### 3.2 In-vitro experiments: Leaflet geometry

Three Live-AVVs were constructed for each of five leaflet geometries: *ℓ*_*v*_*/d* = 0.625, 0.75, 0.875, 1.0, 1.25. For each Live-AVV, six dimensionless chordae lengths *ℓ*_*c*_*/*(*L* − *ℓ*_*v*_) were evaluated. Live-AVVs were secured using a custom built holder that enabled mounting within the pulse duplicator (Supplementary Information). The minimum chordae length, or reference length, was established by applying a standardized tensile force of 4 N to the exposed chordae strands. This force approximates physiological loading conditions across the mitral subvalvular apparatus as reported in literature, corresponding to an average force of approximately 0.5 N per individual chordae [46–49]. Chordae length was tracked using the exposed strand length, defined as the portion of chordae strand external to the holder (Supplementary Information). The remaining segment was contained within the holder. Therefore, decreasing the exposed length increased the effective internal chordae length. To define the longest chordae length, the exposed length was reduced by 2.0 cm from the reference length by retracting the strands back into the holder. The Live-AVV was then mounted onto the pulse duplicator and chordae length was varied by incrementally increasing the exposed length, thereby shortening the internal chordae length. Exposed length was increased stepwise by +0.5 cm, +0.5 cm, +0.5 cm, +0.25 cm and +0.25 cm, with testing performed at each condition.

### 3.3 In-vitro experiments: Fixed design parameters

For initial in-vitro experiments, the following parameters were kept fixed: *α* = 0.9, *d* = 14.4 mm, L = 47 mm, and *ℓ*_L_ = 20 mm. The value of *α* was chosen to approximate physiological AVV geometry [50, 51]. The value of *d* corresponds to an estimated mitral annulus size of a 2 year old child. Conventional options for pediatric AVV replacement, including mechanical and bioprosthetic prostheses, are largely limited to annulus sizes greater than 19 mm [4, 6–8]. The selected annulus dimension therefore represents a clinically relevant pediatric population for whom currently available prosthetic options remain limited. In addition, this annulus size was compatible with available donor valve tissue dimensions. Larger annulus geometries required composite leaflet construction due to insufficient donor tissue size, introducing geometric variability. The selected dimensions permitted consistent fabrication of single-piece leaflets across experiments. Other Live-AVV dimensions were extrapolated from these fixed parameters. Leaflet length (*ℓ*_*v*_) variation is detailed above. Leaflet width is the semiperimeter of the elliptical annulus. We added 2 mm width on each side to ensure sufficient tissue for commissural suturing and to prevent narrowing of the commissural region during annulus attachment. Leaflet geometry and measurement steps are detailed in the Supplementary Information.

### 3.4 In-vitro experiments: Fabrication of chordae tendineae

Artificial chordae tendineae were constructed from expanded polytetrafluoroethylene (ePTFE) sutures (GORE-TEX Suture, CV-7, TTc-13). Detailed steps of chordae tendineae fabrication can be found in the Supplementary Information. Chordal loops were created using Hegar dilators to ensure reproducible loop length.

### 3.5 In-vitro experiments: Fabrication of Live-AVV

Prepared leaflets were sutured onto an elliptical polymer annulus measuring 14.4 mm × 16 mm, fabricated from polyvinyl siloxane. Two identical leaflets were secured to the annulus to establish a symmetric bi-leaflet configuration. Commissural regions were reinforced, followed by attachment of ePTFE neochordae along the leaflet free edge. Detailed fabrication steps, including annulus suturing strategy, commissure formation, and chordal attachment methodology can be found in the Supplementary Information.

### 3.6 In-vitro experiments: Static testing

Completed Live-AVV constructs first underwent static evaluation to assess gross leaflet competence. Valves were mounted onto a custom built holder (Supplementary Information) that secured the polymer annulus while allowing controlled low-pressure water flow across the Live-AVV. This configuration enabled direct visualization of the atrial surface during static pressurization. Under low hydrostatic loading conditions, leaflets were assessed qualitatively for evidence of regurgitant leakage and leaflet prolapse. Regions demonstrating focal leakage were identified and corrected via targeted leaflet repair. Of those valves requiring repair, repair strategies fell into two categories: 1) placement of an additional slit stitch to reinforce commissural closure and prevent low volume focal leakage and 2) plication of the lateral edge of the leaflet at the commissure to better align with the opposing leaflet, aiming to correct small discrepancies in leaflet height. Live-AVVs were statically re-tested following revision and repaired as needed until minimal to no regurgitant leakage was observed. Following the construction of the initial 15 Live-AVVs, these two repair categories were preemptively incorporated into the standard technique, substantially reducing the need for repair during static testing in future Live-AVVs used in the additional in-vitro and in-vivo experiments.

### 3.7 In-vitro experiments: Dynamic testing

Live-AVV constructs that demonstrated minimal to no visible regurgitant leakage under static assessment were subsequently evaluated in a pulse duplicator system (Supplementary Information). The ViVitro platform is a cardiac pulse duplicator that reproduces physiological left heart flow and pressure waveforms through programmable control of heart rate, systolic duration, cardiac output, and afterload. Pulsatile flow conditions were selected according to ISO guidelines for toddler-aged left-heart conditions, in order to approximate physiological hemodynamics corresponding to an annulus size of 14.4 mm × 16 mm. Test parameters were as follows: systolic duration of 45%, mean arterial pressure of 65 mmHg, heart rate of 100 bpm, and cardiac output of 1.5 L/min. During testing, flow profiles and transvalvular pressure gradients were recorded across each cardiac cycle. Hydrodynamic performance parameters, including total regurgitant fraction and mean transvalvular gradient, were calculated according to established pulse duplicator protocols. Simultaneously, echocardiographic imaging was performed through the custom built holder to visualize leaflet motion. Coaptation and billow height were quantified from these images. Coaptation height was defined as the distance from the leaflet free edge to the highest point of apposition between the two leaflets during systole. Billow height was defined as the maximum systolic displacement of the leaflet into the atrium, measured as the perpendicular distance from the plane connecting the two leaflet-annulus hinge points to the highest point of the leaflet belly.

### 3.8 In-vivo experiments: Study design

A sample size of *n* = 3 was used to demonstrate preliminary in-vivo feasibility and functional performance of the optimized Live-AVV in a porcine model. Live-AVV performance was assessed through comprehensive intraoperative monitoring, including echocardiography to evaluate regurgitation, leaflet coaptation and transvalvular pressure gradients. Invasive hemodynamic measurements obtained from right atrial and right ventricular pressure lines were used to corroborate gradient assessment.

All procedures involving human-derived materials were conducted in accordance with relevant institutional guidelines and regulations. Procurement and use of human donor tissues were approved by the Institutional Review Board at Boston Children’s Hospital (IRB-P000045594). All human tissue was obtained through partnership with a registered local organ procurement organization, with written informed consent for research use obtained from the donor or the donor’s legal next of kin. We are grateful to the donor families for their generosity, which made this work possible and enables advancement of research in their loved ones’ honor. Handling of human-derived materials and associated biosafety procedures were reviewed and approved by the Institutional Biosafety Committee of Boston Children’s Hospital (IBC-P00001953). All animal experiments were conducted in accordance with institutional guidelines and regulations and were approved by the Institutional Animal Care and Use Committee at Boston Children’s Hospital (IACUC-P00002762). Key portions of the operations were performed by experienced cardiac surgeons.

Two Live-AVV configurations were attempted for each animal experiment. One configuration consisted of a donor valve sutured onto a polymer annulus prepared the day prior to surgery (ring configuration). Annulus dimensions were estimated according to recipient animal body weight (6-8 kg), as direct measurements were not available prior to the day of the procedure. Live-AVV construction followed the same standardized protocol used for in-vitro experiments, described in detail in the Supplementary Information. Leaflet geometry was defined by optimized dimensionless geometric parameters identified from in-vitro testing: *ℓ*_*v*_*/d* = 1.1, *ℓ*_*c*_*/*(*L* − *ℓ*_*v*_) = 1.1. The second configuration consisted of a Live-AVV constructed intraoperatively and sutured directly to the native annulus (no ring configuration). Live-AVV dimensions were derived from direct measurements of the native tricuspid annulus diameter (*d*) and annulus-to-papillary attachment distance (*L*) as determined by the surgeon based on ventricular geometry. Live-AVV dimensions were calculated as follows: leaflet free edge width (mm) *ℓ* = (*ℓ /d*)(*d*), leaflet annular edge width 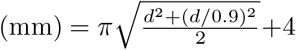 and 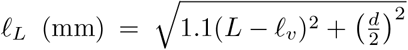. Due to the larger annulus size, composite leaflet construction was required in the third animal experiment. Two donor leaflets were joined using simple interrupted sutures, followed by standard leaflet measurement and trimming according to the standardized in-vitro protocol. Next, artificial chordae tendineae were attached to the leaflet free edge followed by implantation into the recipient heart.

### 3.9 In-vivo experiments: Recipient operation

Recipient animals were premedicated with intramuscular tiletamine-zolazepam, xylazine, and atropine sulfate. General anesthesia was then induced using inhaled isoflurane and intravenous lines were placed. The carotid artery and bilateral external jugular veins were exposed and cannulated for intraoperative hemodynamic monitoring. A median sternotomy was performed to access the mediastinum and the pericardium was opened to expose the heart. The recipient was systemically heparinized and cardiopulmonary bypass (CPB) initiated using standard aortic and bicaval venous cannulation. Animals were cooled to 32°C during CPB. A right atriotomy was performed to expose the native tricuspid valve and *d* was measured. After assessing native ventricle geometry, a target *L* was determined. Next, the native tricuspid valve leaflets and papillary heads were resected. The Live-AVV was then implanted at the native annulus position. The chordae loops were anchored at their convergence point to the ventricular myocardium at *L* distance from the annulus via transmyocardial attachment through the free wall of the ventricle with pledgets secured on the epicardium to distribute tension and prevent myocardial tearing. Future studies will incorporate fixation of chordae loops to papillary muscle heads rather than transmyocardial attachment to more closely replicate native chordal anatomy. Following implantation, animals were rewarmed to 37°C and weaned off CPB. Live-AVV performance was evaluated off CPB using intraoperative echocardiography, including two-dimensional imaging and Doppler assessment. Following completion of imaging, the animal was re-cooled to 32°C and CPB was reinitiated. The first Live-AVV configuration was then explanted, and a second configuration was implanted using the same surgical technique described above. The animal was subsequently rewarmed to 37°C and weaned from CPB for repeat echocardiographic evaluation. In the first two animal studies, the ring configuration was implanted first, followed by the no ring configuration. To minimize potential confounding from the second bypass run on valve performance, the order of configurations was reversed in the third animal. Upon completion of data acquisition, animals were euthanized by exsanguination in accordance with institutional animal care protocols.

### 3.10 LLM usage

We used GPT 5.1 for generating parts of the data analysis code that was used to perform the analysis and optimization of the experimental data.

## Supplementary information

Please see accompanying supplementary files for additional details.

## Acknowledgments

We acknowledge Nicholas Kneier and Shannen Kizilski for their help in setting up the Vivitro experiments. We thank Mengfei He for his guidance on the polymeric valve experiments. We thank Ana Vargas for her assistance in biaxial testing analysis and Vivian Nguyen for her assistance with the in-vivo surgeries. We also acknowledge Kelsey Navin and Arthur Nedder for their assistance with perfusion and animal facility support, respectively. We further acknowledge the contributions of heart donors and their families, as well as the United Network for Organ Sharing (UNOS) and affiliated organ procurement organizations, without whom this work would not be possible.

## Author information

These authors contributed equally: Paul R. Kaneelil, Katrina J. Hon

### Contributions

S.E., L.M., P.R.K., and K.J.H. conceived the project; P.R.K. performed the polymeric valve experiments and designed the geometric valve design with input from L.M; K.J.H. and D.P.R. performed the in-vitro experiments with input from P.R.K; K.J.H, D.P.R., N.T., and S.E. performed the in-vivo experiments and N.T. analyzed and interpreted the echocardiography data; P.R.K. and K.J.H. analyzed data and wrote the first draft; All co-authors edited the paper.

## Supplementary Information

### I. POLYMER VALVE EXPERIMENTS

The prototyping experiments with the conical polymeric valves were done in a custom built flow chamber. See Ref. [1] for more details. The valves had a half cone angle of *β* = 10^*o*^ and a base annulus diameter of *d* = 24 mm. The leaflet length *ℓ*_*v*_, slit length *ℓ*_*s*_, and chordae length *ℓ*_*c*_ were all varied. Four silk chordae were embedded in the polymer while casting the valve, as shown in Fig. S1(a). Figure S1(b) and (c) show side view images of the valve and Fig. S1(d) and (e) show surgeon’s view images of the valve in diastole and systole, respectively. During an experiment, we prescribe the flow field with a pressure pump. The pressure profile is set to ramp linearly over a period of time. The pressure-flow rate data (*p* − *Q*) from our systole experiments, where the flow is in the diverging direction of the cone, is shown in Fig. S1(f). The data shows that the closing dynamics of the valve is independent of chordae length *ℓ*_*c*_*/*(*L* − *ℓ*_*v*_) and slit length *ℓ*_*s*_*/ℓ*_*v*_ and is only a function of the valve length *ℓ*_*v*_*/d*. Note that the closing dynamics shown by the *p Q* data will correspond to the early systolic regurgitation through the valve, which represents the backflow as the leaflets are closing. Figure S1(g) shows the regurgitant flow rate *Q* as a function of time for various different rescaled chordae length 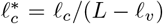. The elbow where the flow rate sharply decreases corresponds to coaptation of the leaflets. The data shows that there is a range of intermediate *ℓ*_*c*_*/*(*L* − *ℓ*_*v*_) that minimizes regurgitant flow through the coapted leaflets, which we refer to as late systolic regurgitation. Taken together, the important parameters that control systolic function are *ℓ*_*v*_*/d* and *ℓ*_*c*_*/*(*L* − *ℓ*_*v*_). Figure S1(h) shows the *p* − *Q* data for diastole when varying the rescaled slit length *ℓ*_*s*_*/ℓ*_*v*_. We see that a higher slit length leads to lower resistance in flow. To minimize the risk of stenosis, we choose to maximize the slit length as much as possible. Details regarding how this is handled for the Live-AVV are given in Fig. S4.

**FIG. S1.**
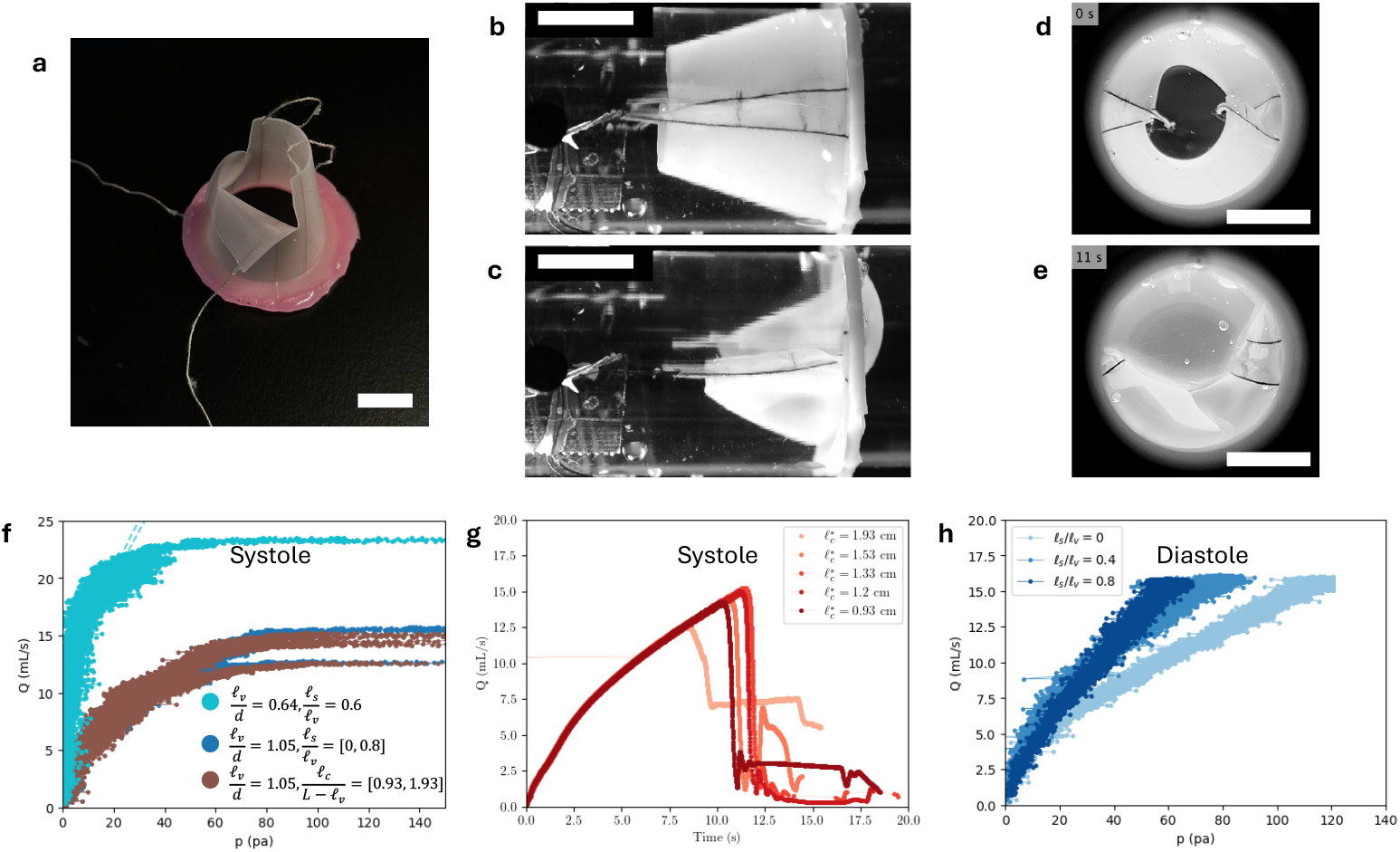
Experiments with a polymer valve. (a) Image showing a polymer valve of cone angle 10^*o*^ with two pairs of silk chordae embedded in the polymer. (b-c) Side view images of the valve in open and coapted positions, respectively. (d-e) Surgeon’s view images of the valve in open and coapted positions, respectively. (f) Pressure-flow rate (*p* − *Q*) curve of the polymer valve during systole for various configurations. (g) Flow rate over time plot for various dimensionless chordae length 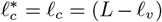. (h) Pressure-flow rate curve of the polymer valve during diastole for valves with different slit lengths. All scale bars represent 10 mm.

### II. VALVE CONSTRUCTION AND TESTING

**FIG. S2.**
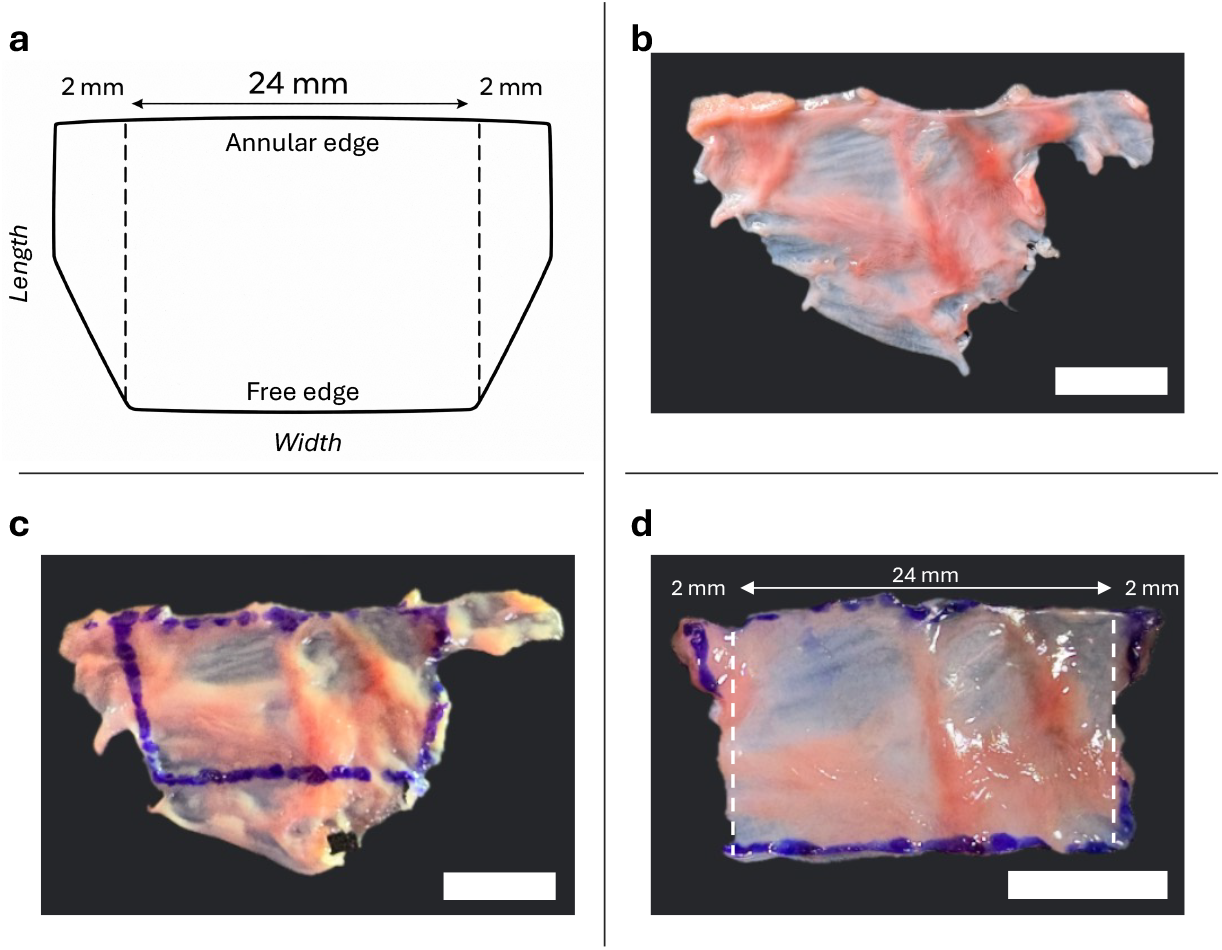
Leaflet geometry and trimming. (a) Schematic of leaflet geometry showing central width and 2 mm extra width on each side of annular edge. (b) Dissected native tricuspid leaflet. (c) Leaflet with marked trimming boundaries. (d) Final trimmed leaflet. All scale bars represent 10 mm.

**FIG. S3.**
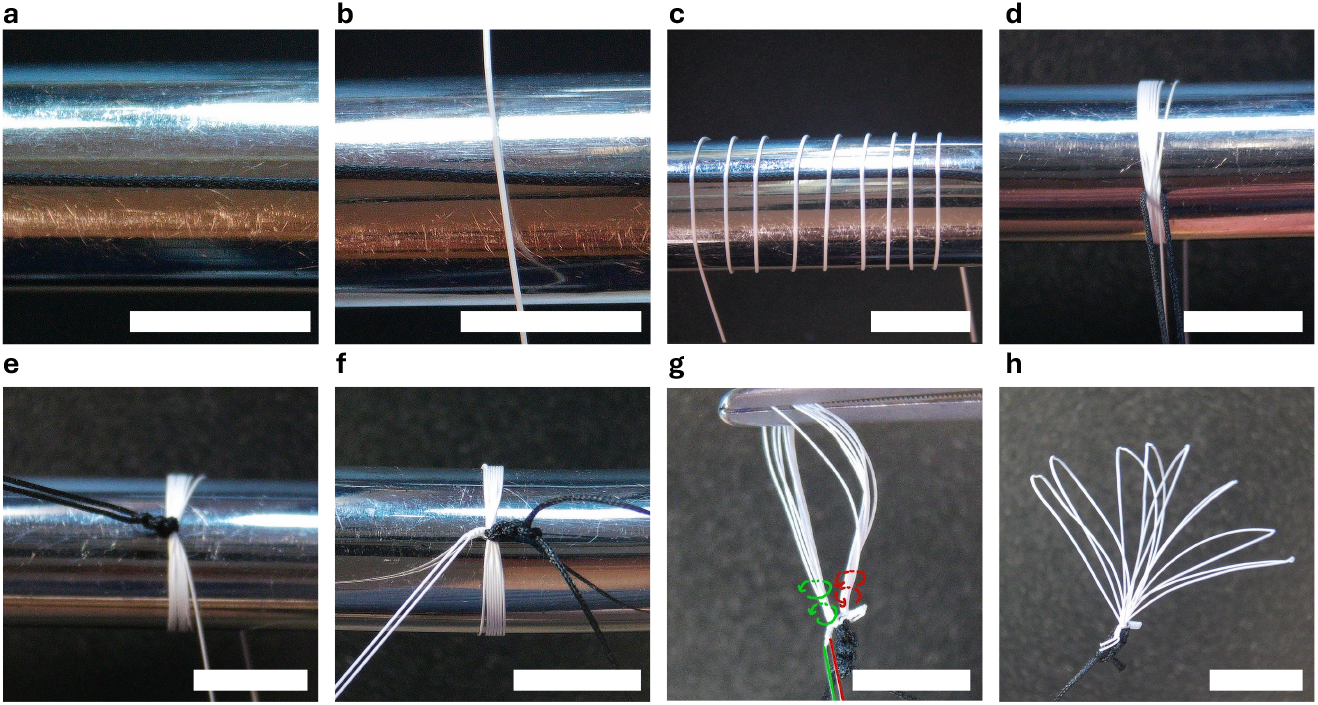
Artificial chordae tendineae construction. (a) Align silk suture along a Hegar dilator. (b) Position ePTFE (GORE-TEX Suture, CV-7, TTc-13) suture perpendicular over silk. (c) Wrap ePTFE total of eight times (one for each loop) around silk and Hegar. (d) Consolidate loops. (e) Knot silk. (f) Knot ePTFE. (g) Pass ePTFE strands twice through loops in opposite directions (i.e., one strand towards (red) and one strand away (green) from operator) and knot both strands together after. (h). Trim excess ePTFE. Repeat steps for total of two chorda tendinea bundles per valve. All scale bars represent 10 mm.

**FIG. S4.**
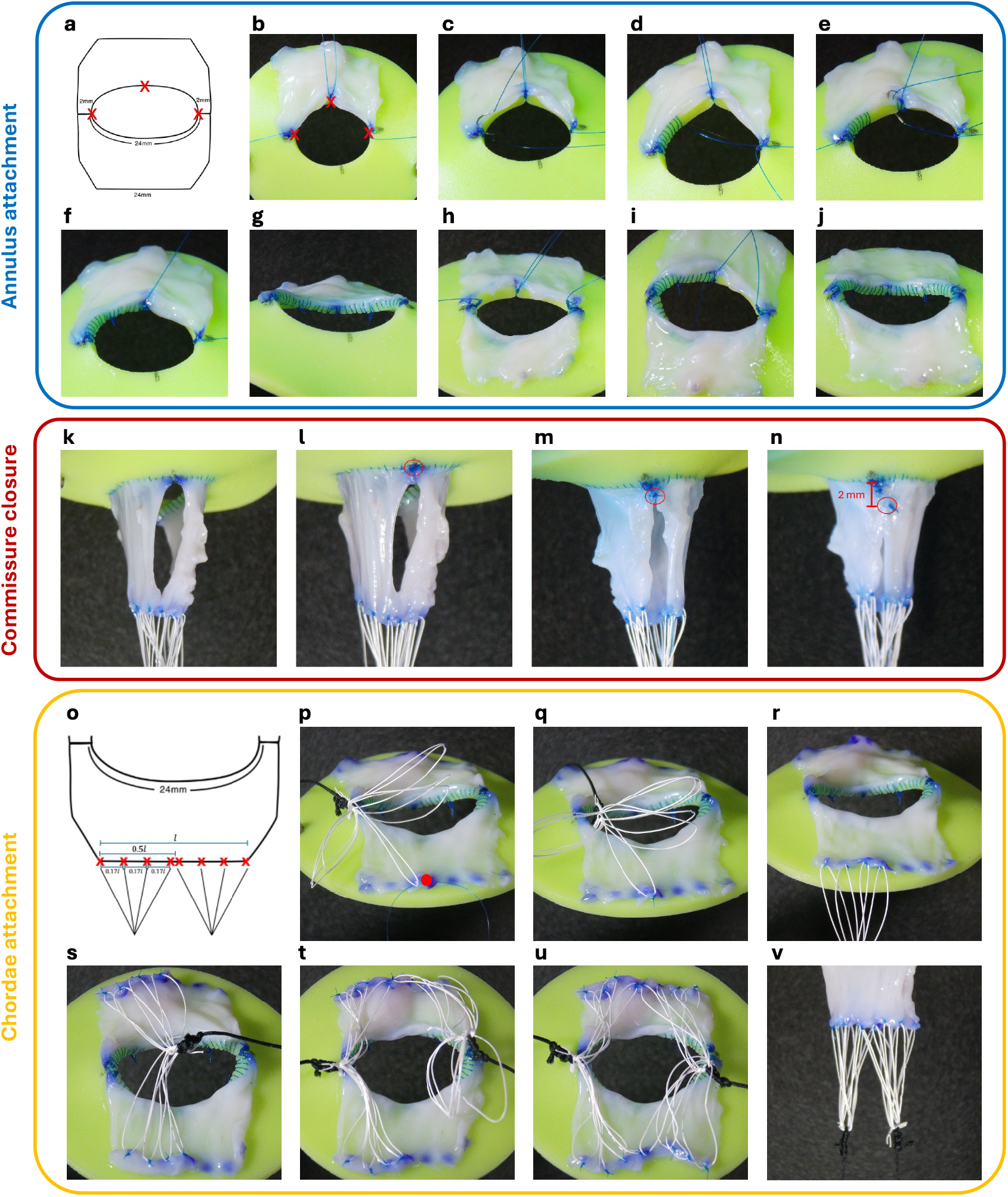
Live-AVV construction. (a-j) Annulus attachment. (a) Schematic of leaflet geometry with initial tagging locations (red X), leaving 2 mm “wings” at the annular edge. (b) Leaflet positioned and tagged on annulus. (c-d) Running suture from edge to midpoint, stopping at one-quarter circumference. (e-f) Running suture from midpoint to meet opposing suture. (g) Remaining quarter completed. (h-j) Second leaflet tagged, sutured, and secured to complete annulus attachment. (k-o) Commissure closure. (k) Side profile prior to closure. (l) Annulus stitch placed (red circle). (m) “Wing” stitch placed to approximate annular edge corners (red circle). (n) Slit stitch placed 2 mm from annulus (red circle). (o-v) Chordae attachment. (o) Schematic of chordae attachment sites along the leaflet free edge. (p-q) Central chordae attached and secured adjacent to midpoint (red dot). (r) Additional chordae secured along same side. (t) Remaining four chordae attached to contralateral side. (t-u) Process repeated for second chordae bundle. (v) Completed valve with chordae. For scale, the elliptical annulus opening has a major diameter of 16 mm and minor diameter 14.4 mm, and the leaflet length is 18 mm. This leaflet length is an illustrative example rather than the optimal construct.

**FIG. S5.**
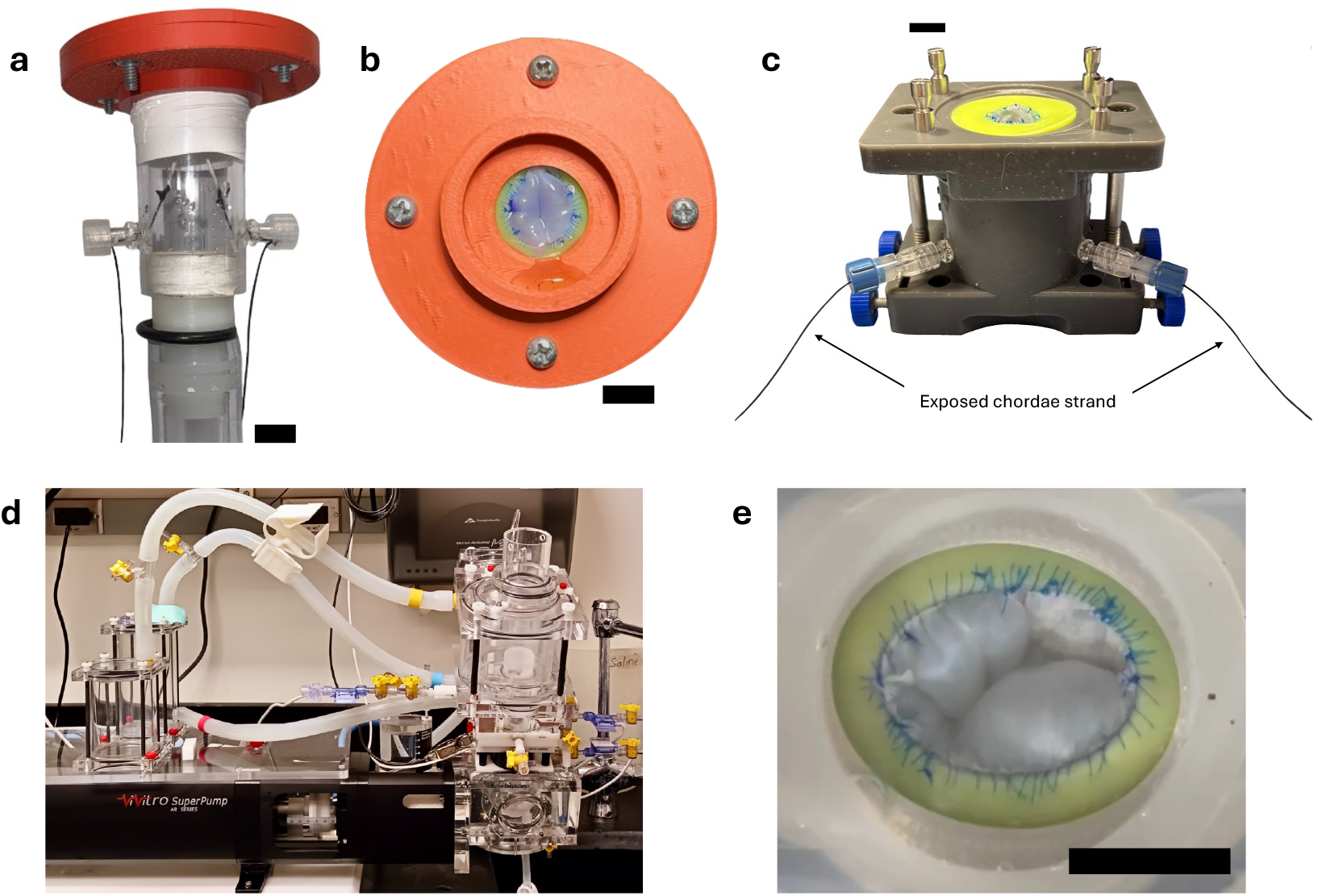
In-vitro Live-AVV testing. (a) Custom built air-water interface tester. (b) Surgeon’s view of mounted valve in air-water interface tester. (c) Custom built Live-AVV holder for ViVitro system, including exposed chordae strands used to determine chordae length. (d) ViVitro pulse duplicator system. (e) Surgeon’s view of mounted valve in ViVitro system. All scale bars represent 10 mm.

### III. EXTENDED IN-VITRO DATA

Figure S6 shows the extended data from the in-vitro experiments, including regurgitation, coaptation height, and billow height data. To quantify the relationship between *R, d*_*c*_*/ℓ*_*v*_, and *d*_*b*_*/d*, as shown in Fig. 2(f), we performed an exponential regression of the form log(*R*) ~ *β*_1_(*d*_*c*_*/ℓ*_*v*_)+ *β*_2_(*d*_*b*_*/d*)+ *β*_3_(*d*_*c*_*/ℓ*_*v*_)^2^ + *β*_4_(*d*_*b*_*/d*)^2^ + *β*_5_(*d*_*c*_*/ℓ*_*v*_)(*d*_*b*_*/d*). The constants from that regression are as follows: *β*_1_ = *-*10.9 *±* 2.54 (*p*< 0.001), *β*_2_ = *-*11.93 *±* 3.27 (*p* = 0.001), *β*_3_ = 7.82 *±* 1.93 (*p*< 0.001), *β*_4_ = 17.86 *±* 3.63 (*p*< 0.001), and *β*_5_ = 10.1 *±* 4.29 (*p* = 0.027).

**FIG. S6.**
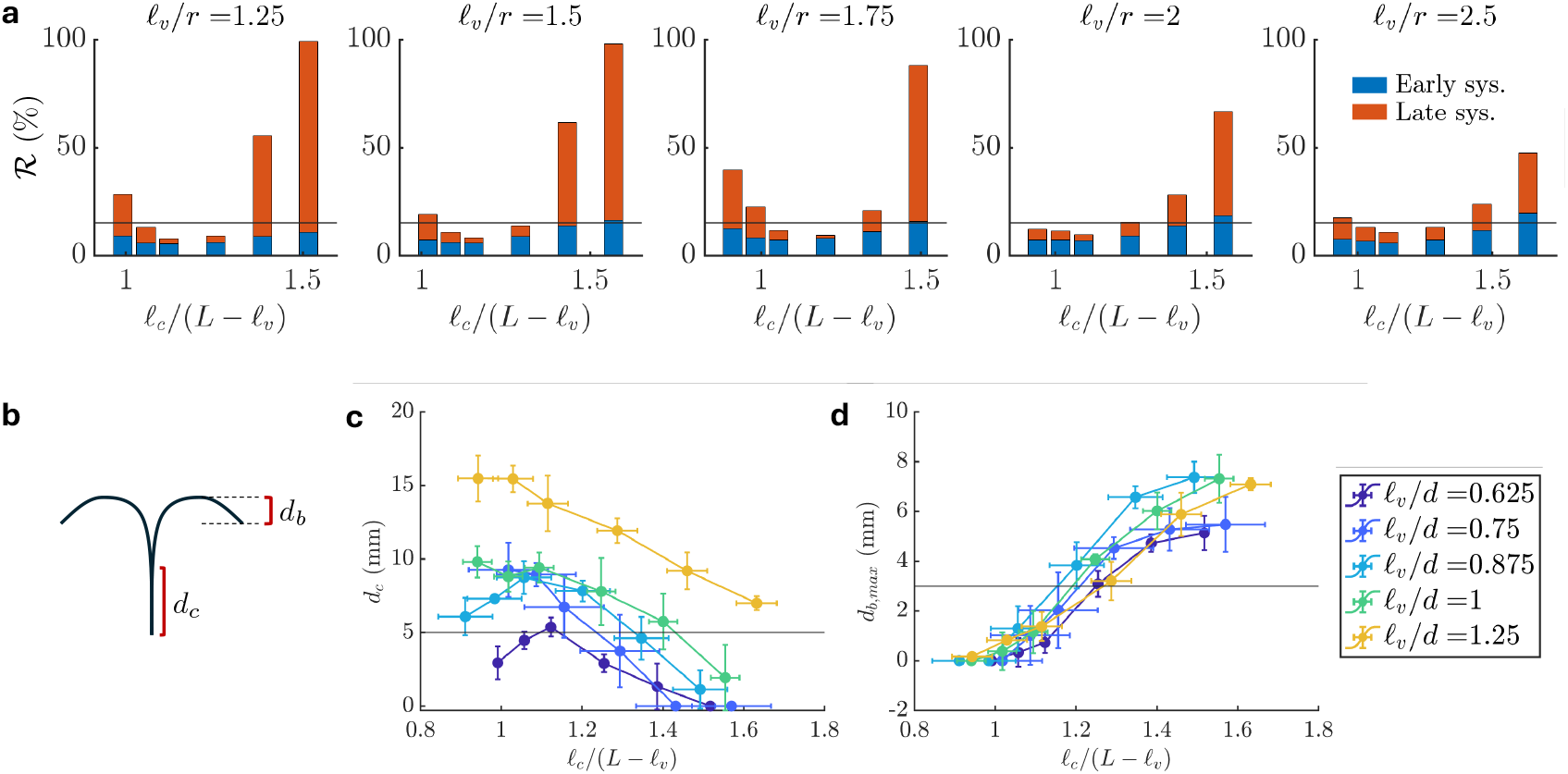
Extended data from in-vitro experiments. (a) Total regurgitant fraction further classified as early systole (blue) and late systole (orange) regurgitation. The former corresponds to the leakage during the closing of the valve and the latter refers to the leakage once the leaflets have coapted. We note that more than half of the leakage is during early systole when *R* is below the clinical threshold (horizontal black line). (b) Schematic showing the definition of coaptation height *d*_*c*_ and billow height *d*_*b*_. (c) Coaptation height *d*_*c*_ as a function of *ℓ*_*c*_*/*(*L* − *ℓ*_*v*_), where the horizontal black line corresponds to the clinical target of 5 mm. (d) Maximum billow height *d*_*b,max*_, defined as the maximum billow of the two leaflets, as a function of *ℓ*_*c*_*/*(*L* − *ℓ*_*v*_), where the horizontal black line corresponds to the clinical target of 3 mm.

### IV. ADDITIONAL IN-VITRO EXPERIMENTS

Additional in-vitro experiments were conducted to evaluate the effects of varying key parameters that were held constant in initial experiments. Namely, we varied loop length *ℓ*_*L*_, slit length, and length from annulus to papillary attachment location *L* (thus *L/d*). In initial experiments, chordae rupture occurred in 40% of Live-AVVs at the shortest chordae length due to excessive tension placed on the leaflet free edge; this condition was therefore excluded from this round of testing.

We found that loop lengths *ℓ*_*L*_ ≥ *d/*2 demonstrated comparable regurgitant fractions and transvalvular pressures, whereas *ℓ*_*L*_ < *d/*2, despite maintaining similar regurgitant fractions, demonstrated increased transvalvular pressures. Reducing slit length by half did not significantly alter regurgitant fraction under conditions where valves already met the clinical target of < 15%. However, at extreme chordae lengths, shorter slit length reduced regurgitant fraction by approximately 50-60%. Notably, Live-AVVs that had been above the clinical threshold with the original slit length remained above the clinical threshold even after reduction of regurgitant fraction by 50-60%. For Live-AVVs with larger *ℓ*_*v*_*/d*, reduction in slit length did not appreciably affect transvalvular pressure; however, an increase was observed for the smallest *ℓ*_*v*_*/d* = 0.625, likely because the slit length was already minimal. The aspect ratio *L/d* used in initial in-vitro experiments (*L/d* = 3.26) was large compared to what might be anatomically possible in-vivo. We performed additional experiments with lower *L/d* and noticed that it increased the range of *ℓ*_*c*_*/*(*L* − *ℓ*_*v*_) over which ℝ and ti ⟨Δ*p*⟩ met the clinical targets. Therefore, the optimal geometry of *ℓ*_*v*_*/d* and *ℓ*_*c*_*/*(*L* − *ℓ*_*v*_) that we found is a suitable and conservative choice independent of *L/d*.

**FIG. S7.**
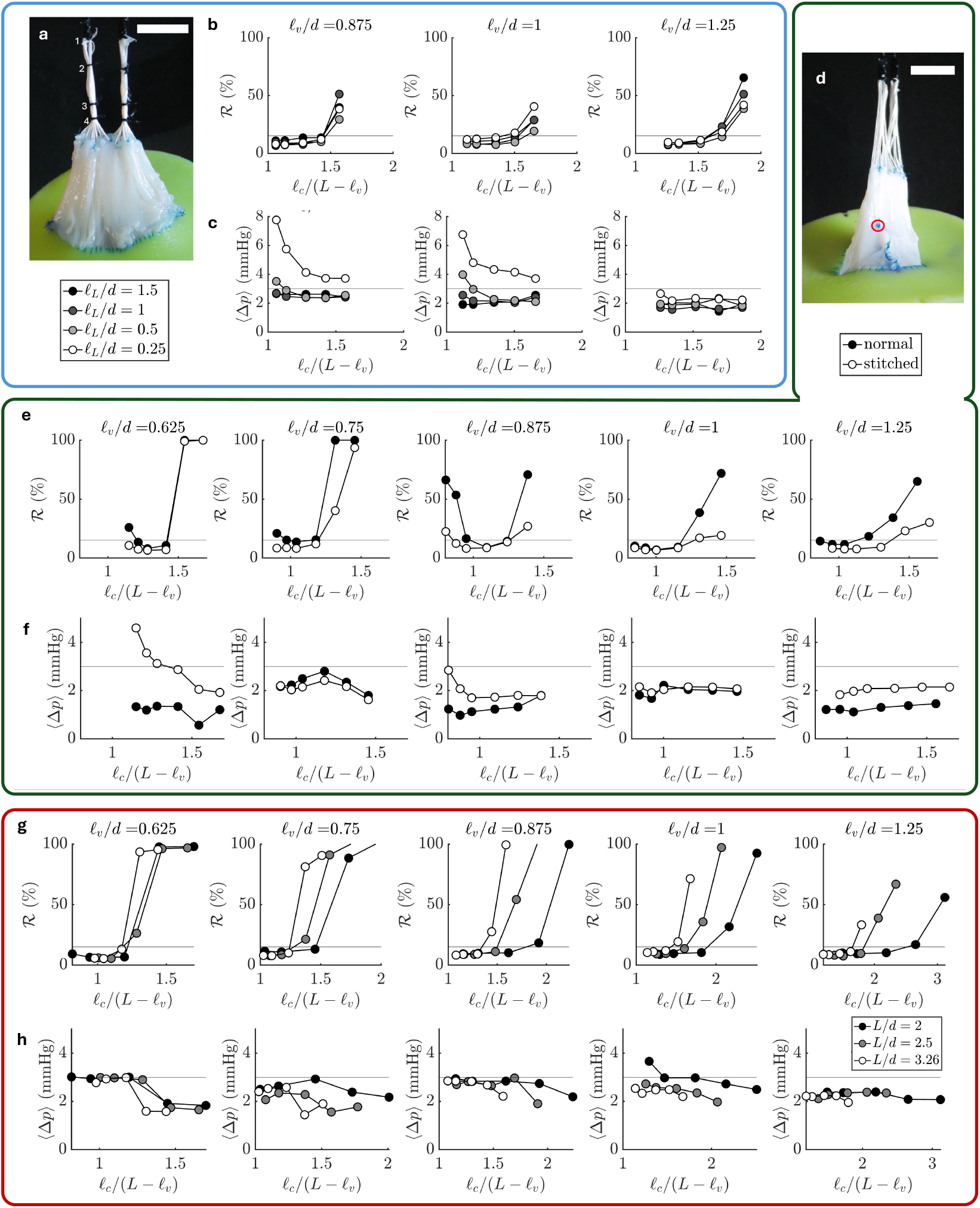
Additional Live-AVV experiments. (a) Representative image of a valve with chordae loop lengths *ℓ*_*L*_*/d* = 1.5 (1), 1 (2), 0.5 (3), and 0.25 (4). Loops were progressively shortened on the same construct by adding silk ties at each specified length, with repeat testing performed after each adjustment, thus testing from longest to shortest *ℓ*_*L*_*/d*. (b-c) *R* and ⟨Δ*p*⟩, respectively, from experiments varying *ℓ*_*L*_*/d* for different *ℓ*_*v*_*/d*. (d) Image of a valve showing the slit stitch (red circle) placed halfway down commissural length to vary the slit length. (e-f) and *p*, respectively, from experiments varying the slit length for different *ℓ*_*v*_*/d*. (g-h) *R* and ⟨Δ*p*⟩, respectively, from experiments varying the aspect ratio *L/d* for different *ℓ*_*v*_*/d*. To vary *L/d*, a custom built holder was redesigned to allow chordae strands to be threaded through three discrete *L* values, allowing us to test *L/d* = 3.26 (original design), 2.5, and 2. Scale bars in (a) and (d) represent 10 mm.

### V. BIAXIAL TESTING

Equibiaxial testing was performed to characterize the anisotropic mechanical properties of mitral valve (MV) and tricuspid valve (TV) tissues and to quantify differences in stretch behavior among individual leaflets. Testing was conducted using a biaxial testing system (Biotester, CellScale, Waterloo, Canada). Specimens were subjected to biaxial loading conditions selected to approximate physiologic stresses experienced by left-heart (MV) and right-heart (TV) valves. Additional details regarding the equibiaxial testing protocol have been described previously [2, 3].

Both MV and TV tissues demonstrated nonlinear, anisotropic mechanical behavior under equibiaxial loading, as shown in Fig. S8. Under representative physiologic loading conditions, MV leaflets stretched about 19% to 25% in the radial direction and 9.6% to 20% in the circumferential direction. Similarly, TV leaflets stretched about 14% to 34% in the radial direction and 8.4% to 12% in the circumferential direction.

Since our optimization is based on experimental results, the stretching of the leaflets is accounted for implicitly. The variability in stretching is then the important aspect to consider. At the membrane tensions *T*_MV_ and *T*_TV_ that the mitral and tricuspid leaflets are expected to sustain for a patient approximately 2 years of age (dashed horizontal lines in Fig. S8), the standard deviation in the radial direction for MV leaflets was 3.2% and for TV leaflets was 10%. This variability is essentially captured by the physiological variation and the associated safety factor we have considered for finding the optimal geometry.

**FIG. S8.**
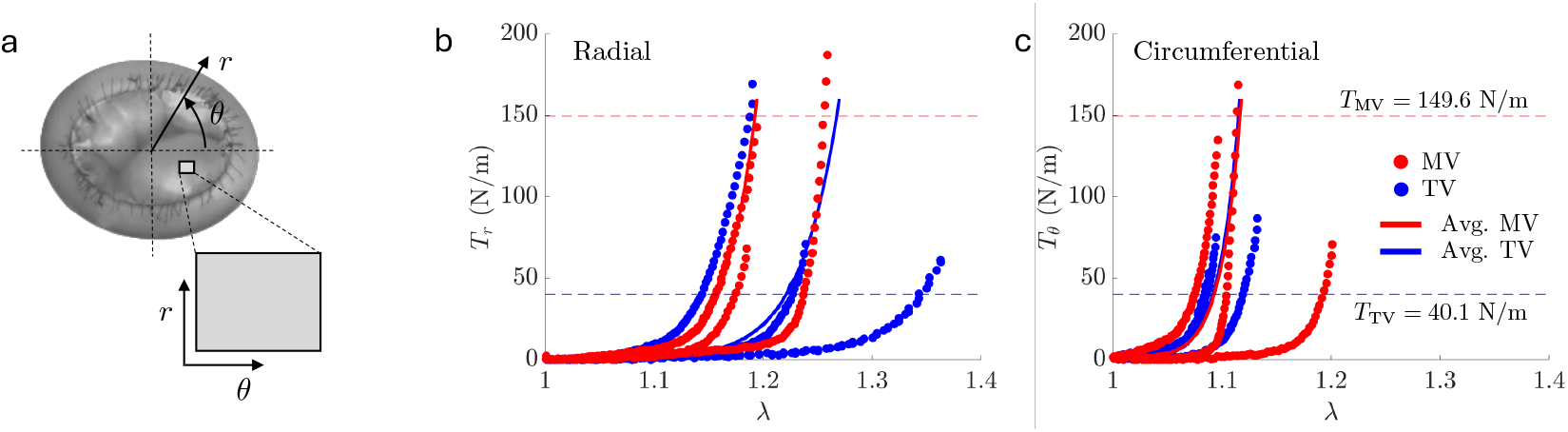
Biaxial testing results for mitral (MV) and tricuspid (TV) leaflet tissue. (a) Schematic showing the radial *r* and circumferential *θ* directions of a valve tissue sample (rectangular) that is used for biaxial testing. (b) Radial membrane tension *T*_*r*_ as a function of stretch λ for MV (red) and TV (blue) samples. The solid line shows the average values calculated using the common stretch method. The horizontal lines represent the expected membrane tensions that the MV and TV leaflets sustain for a patient of 2.3 yrs. (c) Circumferential membrane tension *T*_*θ*_ as a function of λ. Datasets shown in the same color correspond to different samples and thus illustrate the variability in the results.

